# Foraging ecology drives viral community structure in New Zealand’s aquatic birds

**DOI:** 10.64898/2025.12.09.693119

**Authors:** Stephanie J Waller, Janelle R Wierenga, Lia Heremia, Jessica A Darnley, Isa de Vries, Jeremy Dubrulle, Zoe Robinson, Allison K Miller, Chris N Niebuhr, David S Melville, Rob Schuckard, Phil F Battley, Michelle Wille, Ben Alai, Rosalind Cole, Jamie Cooper, Ursula Ellenberg, Graeme Elliott, James Faulkner, Johannes H. Fischer, Lance Hay, David Houston, Bianca C Keys, Jenny Long, Robin Long, Thomas Mattern, Lou McNutt, Peter Moore, Odin Neil, Jake Osborne, Anne-Sophie Pagé, Kevin A Parker, Mike Perry, Brodie Philp, Kalinka Rexer-Huber, James C Russell, Rachael Sagar, Thor T Ruru, Theo Thompson, Leith Thomson, Joris Tinnemans, Lydia Uddstrom, Te Arawhetu Waipoua, Kath Walker, Edin Whitehead, Chrissy Wickes, Melanie J. Young, David Winter, Kate McInnes, Edward C Holmes, Jemma L Geoghegan

**Author notes:** Corresponding author: Jemma Geoghegan.

## Abstract

Wild migratory birds play a major role in the global spread of viruses, yet the diversity, host range and transmission patterns of viruses harboured by migratory species in Aotearoa/New Zealand remain largely unknown. This knowledge gap is critical given New Zealand’s position along major migratory flyways spanning Oceania, Antarctica and east Asia, where understanding viral diversity is key to assessing the risk of viral introductions such as highly pathogenic avian influenza virus and viral dispersal across these regions. To address this, we conducted the first large-scale metatranscriptomic survey of wild birds from New Zealand and its subantarctic islands, collecting 1,348 samples from 31 host species spanning four avian orders. We identified 118 avian viruses from 17 families, including 107 novel species, greatly expanding our knowledge of avian viral diversity. Viral communities differed significantly by host order and foraging behaviour, with scavenger birds harbouring more diverse viromes than non-scavengers. Although no HPAI subtypes were detected, we recovered a low-pathogenic avian influenza A/H1N9 virus from red knots (*Calidris canutus*) and a divergent tobanivirus from Auckland Island teal (*Anas aucklandica*), the first putative avian member of the *Tobaniviridae*. Notably, we detected 12 mammalian-associated viruses, primarily in scavenger birds, including *Hedgehog hepatovirus*, *Rabbit haemorrhagic disease virus 2*, and sea lion astroviruses, with mammalian host reads confirming their dietary origin. This study establishes the first virome baseline for New Zealand’s migratory birds, highlighting the ecological role of foraging in shaping viral communities and improving regional preparedness for HPAI and other emerging avian pathogens.

## Introduction

Despite Aotearoa/New Zealand’s unique avifauna and its position along major migratory flyways(1), little is known about the diversity and ecology of viruses circulating among New Zealand’s wild bird populations. Wild birds host a diverse array of viruses, and the long-distance migration of some species can make them important vectors for viral incursions into new regions, such as the spread of highly pathogenic avian influenza (HPAI) virus subtype H5N1 from Asia to Europe, Americas, Antarctica(1–3), and the spread of Usutu virus from Africa to Europe(4). Previous avian virus investigations in New Zealand have typically only focused on single species(5–7), a specific bird order (8), or an isolated sampling location(9). Few have examined multiple regions or integrated large-scale surveillance approaches(10). In addition, no prior viral surveillance has been conducted in avian populations from New Zealand’s subantarctic islands, despite these islands hosting vast, dense seabird colonies that connect with migratory networks spanning the Southern Ocean(1). This limited knowledge hampers our ability to detect novel or emerging viruses, to identify ecological factors that promote viral transmission, and to assess the potential risks these viruses pose to native wildlife, agriculture and biosecurity. Establishing a comprehensive baseline of viral diversity across New Zealand’s avian communities is therefore critical for detecting and mitigating future disease incursions.

This lack of baseline knowledge has become increasingly concerning given the global spread of HPAI H5N1(11). Since 2020, the H5N1 subclade 2.3.4.4b has driven extensive outbreaks across all continents besides Oceania, causing mass mortality in wild birds, mammals, and poultry, and reshaping global patterns of avian influenza transmission(12). Once primarily associated with only waterfowl(13), recent epizootic waves have demonstrated that migratory seabirds now play a central role in long-distance dissemination, introducing HPAI into new regions, including remote island ecosystems(1,12,14).

Low pathogenic avian influenza viruses have been detected in New Zealand birds for more than two decades(15,16). The only HPAI detection in New Zealand involved a locally adapted subtype (H7N6), which caused limited outbreaks in poultry(17). However, New Zealand’s position within the East Asian–Australasian Flyway, and the connectivity of its subantarctic seabird colonies with Antarctica, make it vulnerable to viral introduction(1). The recent detection of H5N1 in remote island systems such as South Georgia(18), the Falkland Islands(18), and the Crozet and Kerguelen archipelagos(19) heightens the risk of HPAI H5N1 introduction into New Zealand. An incursion into New Zealand could have devastating ecological and economic consequences, threatening native bird species already vulnerable to infectious disease, and imposing severe biosecurity and trade impacts on the poultry and dairy industries (20,21).

Here, we characterise the viral repertoire of wild migratory bird populations across New Zealand and its subantarctic islands. By examining patterns of virus distribution and transmission, this study fills a critical gap in our understanding of avian viral ecology in New Zealand and strengthens preparedness for potential incursions of HPAI and other emerging avian diseases.

## Methods

### Ethics statement

Animal ethics for sampling birds at the Firth of Thames and Motueka Sandspit were obtained from the Massey University Animal Ethics Committee (MUAEC 22/52) and sampling was permitted via the Department of Conservation (Wildlife Act Authority 38111-FAU). All other birds were sampled by the Department of Conservation in accordance with their Wildlife Health Management Standard Operating Procedure.

### Avian oral and cloacal swab sample collection

A total of 690 individual birds were captured using mist nets, cannon netting, or by hand, depending on the species. Oral and cloacal swabs were collected from each bird where possible, totalling 1348 samples. Swabs were then placed into 800ul of RNA stabilisation solution (DNA/RNA Shield, Zymo Research) and were stored at 4°C until the samples were sent to the University of Otago, Dunedin, New Zealand, where they were stored at -80°C until RNA was extracted. More information regarding sample locations, species, swab type and the number of individual birds sampled at each sampling site is provided in Supplementary Table 1.

### Total RNA extraction

Frozen oral and cloacal swabs were defrosted before being placed in ZR BashingBead Lysis Tubes (0.1 mm and 0.5 mm) (Zymo Research) filled with 1 mL of DNA/RNA shield (Zymo Research). Lysis tubes were placed into a mini-beadbeater 24 disruptor (Biospec Products Inc.) and were homogenised for five minutes. Total RNA was extracted using the ZymoBIOMICS MagBead RNA kit (Zymo Research) following the manufacturers protocol. RNA was quantified using a nanodrop. In total, RNA from 683 cloacal and 665 oral swabs were at suitable concentrations for sequencing. Equal volumes of RNA from 1-24 individuals were pooled into 205 groups based on species and location (see Supplementary Table 1).

### RNA sequencing

Extracted RNA was subject to total RNA sequencing. Libraries were prepared using the Illumina Stranded Total RNA Prep with Ribo-Zero Plus (Illumina) and 16 cycles of PCR. Paired-end 150 bp sequencing of the RNA libraries was performed on the Illumina NovaSeq 6000 platform.

### Virome assembly and virus identification

Sequencing reads were quality-trimmed using Trimmomatic v0.38(22). Adaptor sequences were removed, low-quality bases with quality scores of less than 3 were trimmed from the 5′ and 3′ ends, and sliding window trimming (4 bases, average quality <5) was applied.

Reads shorter than 25 nucleotides after trimming were discarded and only paired reads retained after trimming were used for downstream analyses. Following quality control, reads were *de novo* assembled into contigs using MEGAHIT v1.2.9(23). Contigs were then compared against the NCBI nucleotide (nt) and non-redundant protein (nr) databases using BLASTn v2.15.0(24) and DIAMOND v2.1.9(25) to identify viral sequences. A similarity threshold of 1x10⁻⁵ for the nt database and 1x10⁻¹⁰ for the nr database was applied to reduce false positive results.

### Estimating viral transcript abundance estimations

Virus abundance was estimated by mapping reads back to the assembled contigs using Bowtie2 v2.4.4(26) and SAMtools v1.9(27). Sequences representing <0.1% of the read counts in another library and sharing >99% nucleotide identity were considered likely index-hopping artifacts and excluded from further analysis. Total viral abundance estimates for viruses from avian hosts across viral families and orders were compiled across all libraries. Estimated abundances per million reads were standardised to the number of paired reads per library.

### Virus phylogenetic analysis

Translated conserved viral protein sequences (i.e., RNA dependent RNA polymerase (RdRp), DNA polymerase, VP1 or capsid, depending on viral taxonomy) were aligned with representative protein sequences from the same virus family obtained from NCBI RefSeq as well as the closest BLASTp hits using MAFFT v7.490(28). Ambiguously aligned regions were removed using trimAL v1.2rev59(29) with the gap threshold flag set to 0.9.

Phylogenetic trees for each viral species/family/order were then estimated using the maximum likelihood method in IQ-TREE v1.6.12(30), employing the LG amino acid substitution model with 1000 ultra-fast bootstrapping replicates. The resulting phylogenetic trees were annotated using Figtree v1.4.4(31). Phylogenetic comparison with known viruses was used to infer the probable host origin of each virus. Specifically, viruses that were phylogenetically distinct from vertebrate host viruses were assumed to be more likely associated with diet, microbiome or environmental sources. Viruses that clustered with known avian or mammalian viruses were subject to further evolutionary analysis.

### Bipartite network analysis and plots

All plots were created in R v4.3.1 using RStudio v2021.09.1 with the tidyverse ggplot2 package(32). For each library, raw read counts were first standardised as reads per million (RPM), calculated as the number of reads mapping to a viral family divided by the total number of reads in that library, multiplied by one million. To visualise viral community composition, read counts were summarised by sample type (cloacal vs oral swabs) and viral family. The data were then plotted as proportional pie charts using ggplot2(32).

A bipartite network analysis was used to investigate the relationships of viral families or viral species that were or were not shared between avian species. The bipartitie network was created using the ggbipart package(33) within the ggplot2(32) environment. To quantify structural properties of the network, nestedness was calculated using the bipartite package in R(34). Nestedness describes the degree to which species with fewer viral associations harbour subsets of those found in more virus-rich hosts.

To visualise viral community composition, we constructed a heatmap of viral family abundances across libraries. RPM values were normalised by the total RPM within each library so that the values summed to one, allowing direct comparison across samples of differing sequencing depths. Relative viral family abundances were plotted using a heatmap in ggplot2(32). Relative abundances were represented on a log^10^ scale. Libraries were ordered by region and bird order to highlight ecological structure.

To assess which factors (i.e., foraging behaviour, migration behaviour, geographic location, swab type, avian taxonomy) most influenced viral diversity, we compared within-sample (alpha) diversity across these factors. Categories describing foraging and migration behaviours were assigned using information from published literature and consultations with avian experts from the New Zealand Department of Conservation. Foraging status was categorised as: (i) scavenger: a bird species that regularly consumes carrion or scavenges as a major component of their diet, (ii) occasional scavenger: a species that may opportunistically scavenge but does not rely on it, or (iii) non-scavenger: a species not known to scavenge. Migratory status was classified as: (i) migratory (undertaking long-distance seasonal movements), (ii) local migration (short-range or within-country seasonal movements), or (iii) sedentary (remaining in the same region year-round). To determine whether scavenging behaviour was phylogenetically structured, we tested the association between bird order and scavenging status using a contingency table and Fisher’s exact test(35).

Viral diversity was assessed using the Shannon diversity index, calculated from viral family-level relative abundance data (RPM) using the Vegan package(36). Differences in Shannon diversity across each factor were evaluated using a Kruskal–Wallis test. When significant differences were detected, a Dunn’s post hoc test with Benjamini–Hochberg correction for multiple comparisons was applied. Boxplots of Shannon diversity were generated with ggplot2, and statistically significant pairwise comparisons were annotated on the plots using ggpubr.

Similarly, we conducted beta diversity analyses to evaluate whether foraging behaviour, migration behaviour, geographic location, swab type, or avian order influenced overall viral community composition. Beta diversity was assessed using Bray-Curtis dissimilarity, calculated from viral abundance values (RPM). A non-metric multidimensional scaling (NMDS) ordination was then performed (metaMDS, vegan(36)) to visualise differences in viral community composition among samples. Ordination plots were created using ggplot2, with samples coloured by ecological trait or factor, and group ellipses added to illustrate clustering patterns. To statistically test for differences in viral community composition between groups, a Permutational Multivariate Analysis of Variance (PERMANOVA) (adonis2, vegan(36)) was conducted using Bray–Curtis distances. First, the effect of ecological traits and other factors were tested independently. Subsequently, a marginal-effects PERMANOVA including all ecological traits and other factors was performed to determine the relative contribution of each factor to overall community variation. To verify that differences were not driven by heterogeneity in dispersion among groups, beta dispersion was assessed using the betadisper function(36), followed by ANOVA and pairwise permutation tests. Dispersion results were visualised as both centroid plots and boxplots representing the distance of each sample to its group centroid.

To identify viral families significantly associated with different foraging behaviours, an indicator species analysis was performed using the multipatt function in the indicspecies R package(37). This approach identifies taxa characteristic of particular groups by evaluating both their specificity (uniqueness to a group) and fidelity (consistency within a group) through permutation testing (999 permutations). Pairwise differences in abundance between foraging categories (‘non-scavenger’, ‘occasional scavenger’, and ‘scavenger’) were further assessed using non-parametric Wilcoxon rank-sum tests.

### Viral nomenclature

A virus was tentatively considered a novel species if it shared<90% amino acid similarity within the most conserved region (i.e. RdRp/polymerase). For the novel virus sequences identified here we have provided a provisional virus (common) name prior to formal verification by the International Committee on Taxonomy of Viruses (ICTV).

## Results

Overall, 683 cloacal and 665 oropharyngeal swabs were sampled from 31 wild bird species spanning four avian orders (Figure 1a-c, Supplementary Table 1) between November 2023 and March 2024 (i.e., spring to autumn) in New Zealand. These samples were pooled into 205 libraries for total RNA sequencing (Supplementary Table 1). From these data (which comprised a total of 12.5 billion sequencing reads) we identified 17 viral families that were deemed likely to infect birds. Avian viruses from the *Orthoherpesviridae* comprised the largest proportion of viral reads in cloacal swabs, while picornaviruses had the largest proportion of avian viral reads in oropharyngeal swabs.

**Figure 1.**
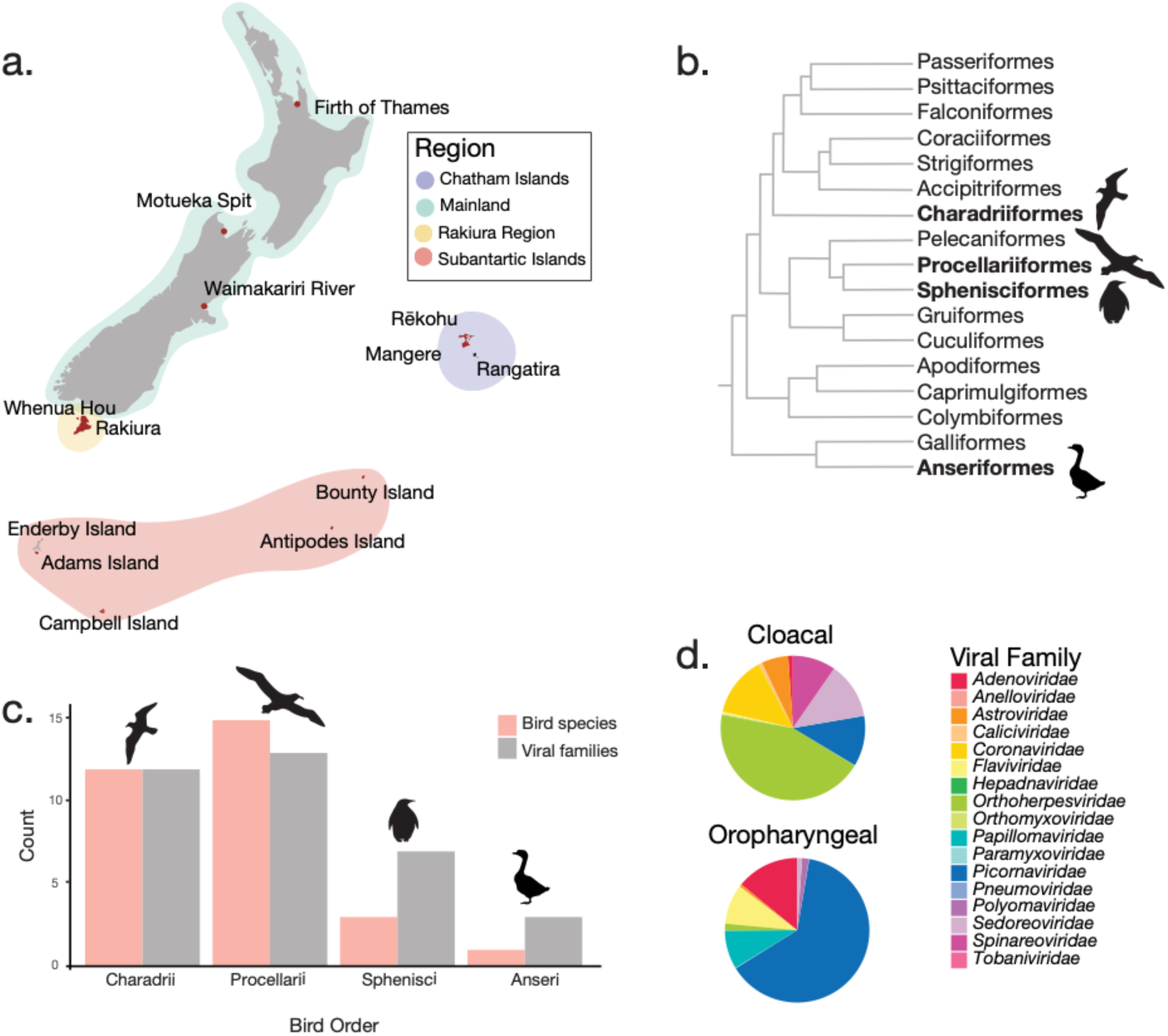
Summary of sampling locations, host diversity, and viral detection. (a) Map of the 13 locations (red) across New Zealand and subantarctic islands where birds were sampled, with the four sampling regions highlighted. (b) Phylogenetic tree showing relationships among 17 avian orders (based on Corfield et al., 2014(38)), with the four sampled orders in bold. (c) The number of bird species sampled and viral families identified within each order. (d) The proportional distribution of viral families detected in cloacal and oral swabs.

Southern black-backed gulls (*Larus dominicanus*) and subantarctic skuas (*Stercorarius antarcticus*) harboured the highest number of viral families (nine and eight, respectively) compared with other species (Figure 2a). In contrast, banded dotterels (*Anarhynchus bicinctus*), red-billed gulls (*Chroicocephalus novaehollandiae*), and common diving petrels (*Pelecanoides urinatrix*) did not harbour any avian viruses. Picornaviruses, papillomaviruses, and orthoherpesviruses were the most widespread viruses, detected in 19, 15, and 13 bird species (of 31), respectively (Figure 2a). In comparison, members of the *Tobaniviridae*, *Polyomaviridae*, *Pneumoviridae*, *Orthomyxoviridae*, and *Anelloviridae* were each identified in only a single avian species (Figure 2a).

**Figure 2.**
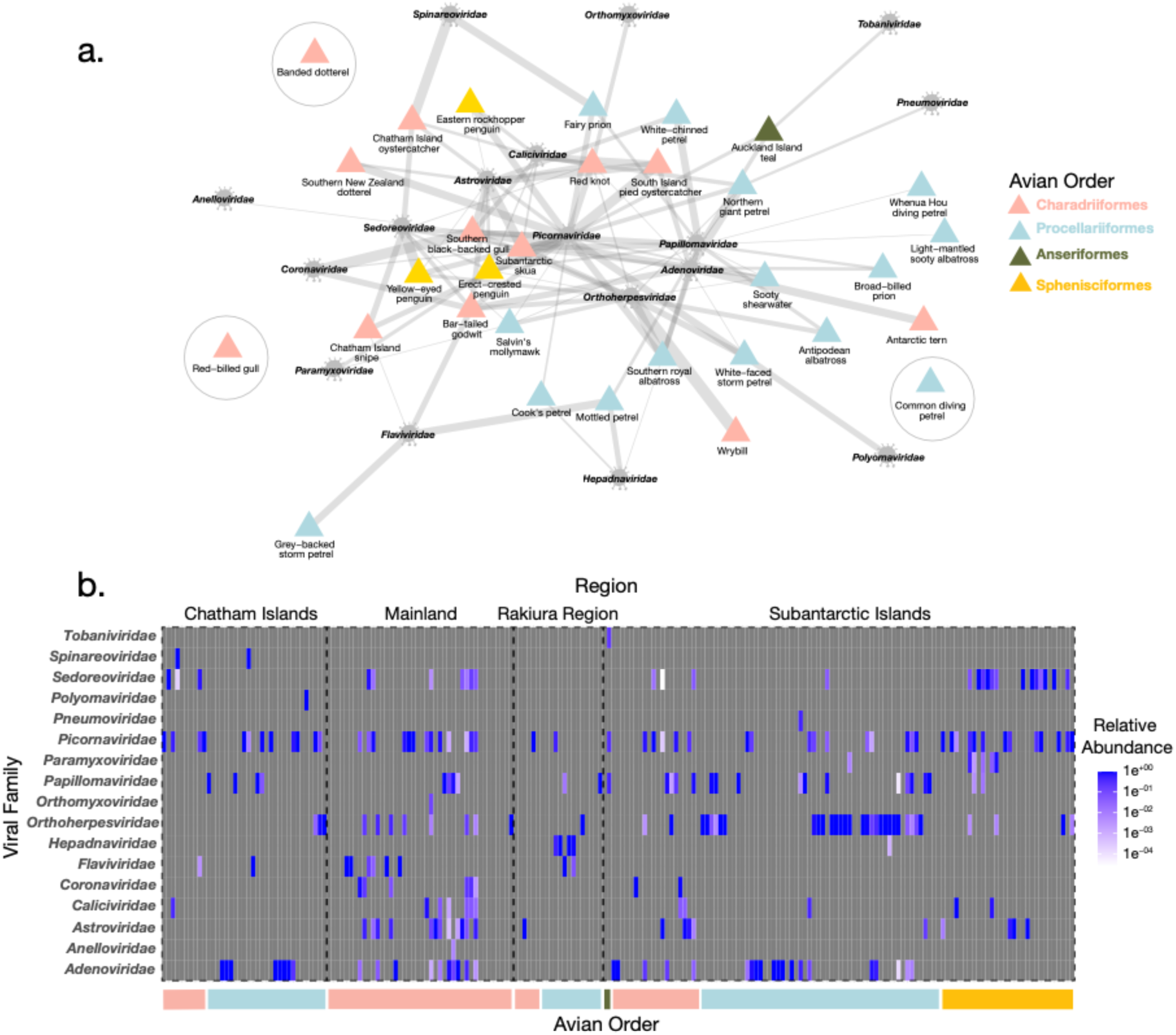
Host–virus associations visualised through a bipartite network and relative abundance heatmap. (a) Bipartite network illustrating associations between sampled bird species and the viral families detected. Each node represents either a bird species, coloured according to avian order, or a viral family, with edges indicating the presence of viral reads in that host species. Edge weights were derived from summed reads per million (RPM) values across libraries for each species. (b) Heatmap showing the relative abundance of viral families in each library, calculated from RPM values normalised by the total RPM per library. Libraries are grouped by region and avian order. Colour intensity indicates relative abundance on a log^10^ scale, ranging from low (white) to high (blue). Coloured bars below the heatmap indicate the avian order of the sampled host species.

Virome composition appeared to be influenced by avian order, with Charadriiformes and Procellariformes exhibiting largely distinct viral associations (Figure 2a). Consistent with this, a bipartite network analysis revealed low nestedness (NODF = 16.75, Figure 2a), indicating that the network was not strongly hierarchical (such that species with fewer detected viruses did not simply harbour subsets of those found in more virus-rich hosts). Rather, the structure appeared more modular or host-specific, suggesting that distinct viral assemblages are associated with different avian taxa. A total of 12 avian viral families were detected in bird samples from the Subantarctic Islands, 11 in mainland New Zealand samples, nine in samples from the Chatham Islands, and six in those from the Rakiura region, although no consistent trend in viral family diversity was apparent across regions or avian orders (Figure 2b).

In total, we identified 107 novel avian viruses and 11 previously described viruses spanning 17 viral families (Figure 3, Figure 4a, Supplementary Table 2). These findings greatly expand the known diversity of avian viruses, particularly within New Zealand’s aquatic birds. Notably, 16 of the viruses detected were closely related (>90% amino acid similarity within the conserved RNA dependent RNA polymerase, ORF1ab, DNA polymerase or L1 region) to those reported in other geographic regions including Australia, Antarctica, Japan, the Falkland Islands, Russia, China, and the United Arab Emirates (Figure 3). Our phylogenetic analysis also revealed that viruses detected in hosts of the same avian order generally clustered together, suggesting host-specific associations (in accordance with the bipartite network analysis), as did viruses sampled from the same geographic region often grouped together, indicating regional structuring of avian viromes (Figure 3).

**Figure 3.**
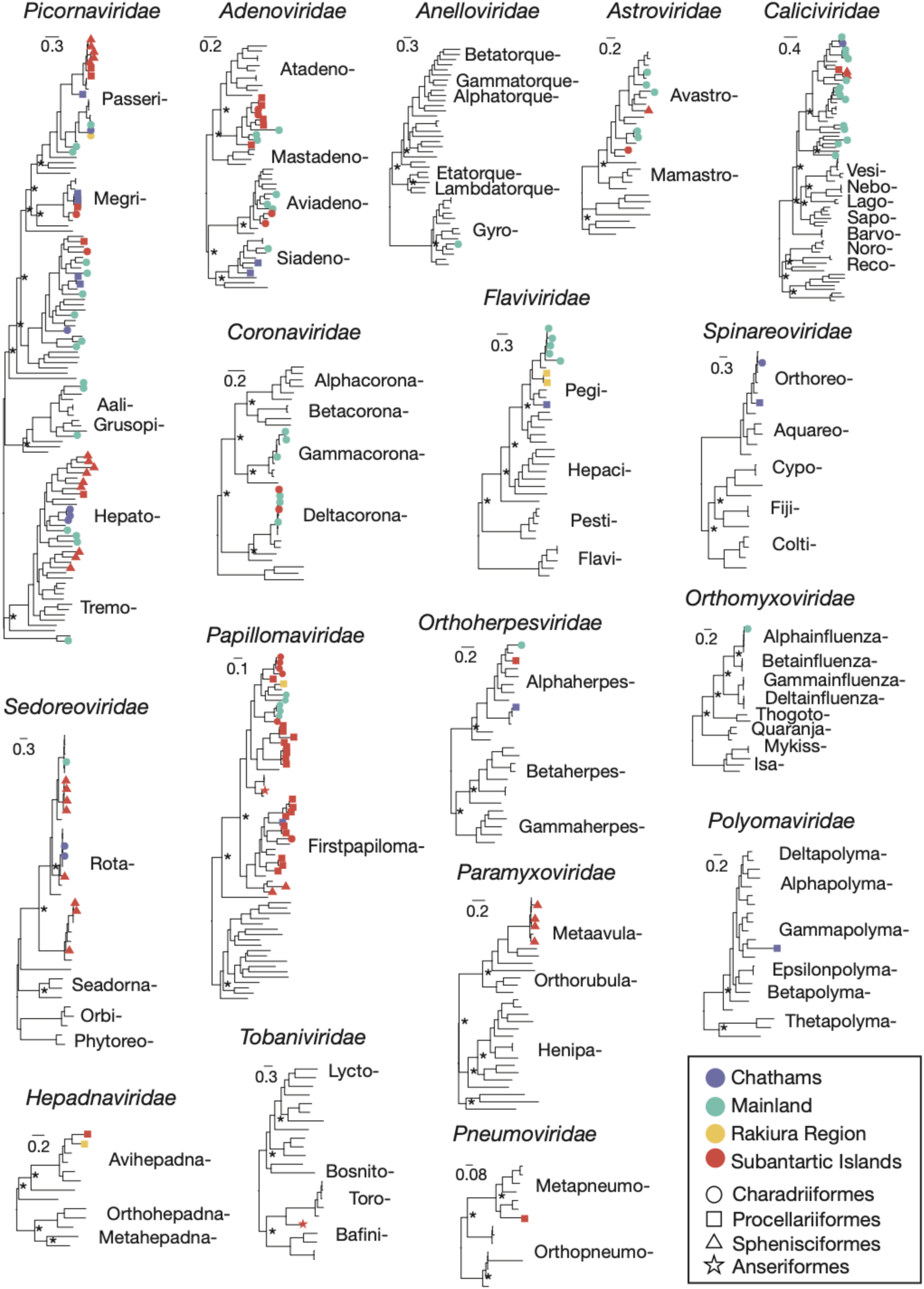
Phylogenetic trees of avian viruses. Maximum likelihood phylogenetic trees were constructed from representative viral transcripts containing RdRp, DNA polymerase, VP1, or capsid sequences from 17 viral families (*Picornaviridae*, *Adenoviridae*, *Anelloviridae*, *Astroviridae*, *Caliciviridae*, *Coronaviridae*, *Flaviviridae*, *Spinareoviridae*, *Sedoreoviridae*, *Papillomaviridae*, *Orthoherpesviridae*, *Orthomyxoviridae*, *Hepadnaviridae*, *Tobaniviridae*, *Paramyxoviridae*, *Pneumoviridae*, *Polyomaviridae*). Viruses identified in this study can be identified by the tip labels, with colours indicating the sampling region and shapes indicating the avian order of the host that was sampled. Branch lengths are proportional to the number of amino acid substitutions per site, and all trees are midpoint-rooted. Main nodes with ultrafast bootstrap values >70% are indicated by an asterisk. Annotated phylogenetic tree files with taxon labels can be found at https://github.com/stephwaller/NZ-Avian-Virome-2023-2024.

**Figure 4.**
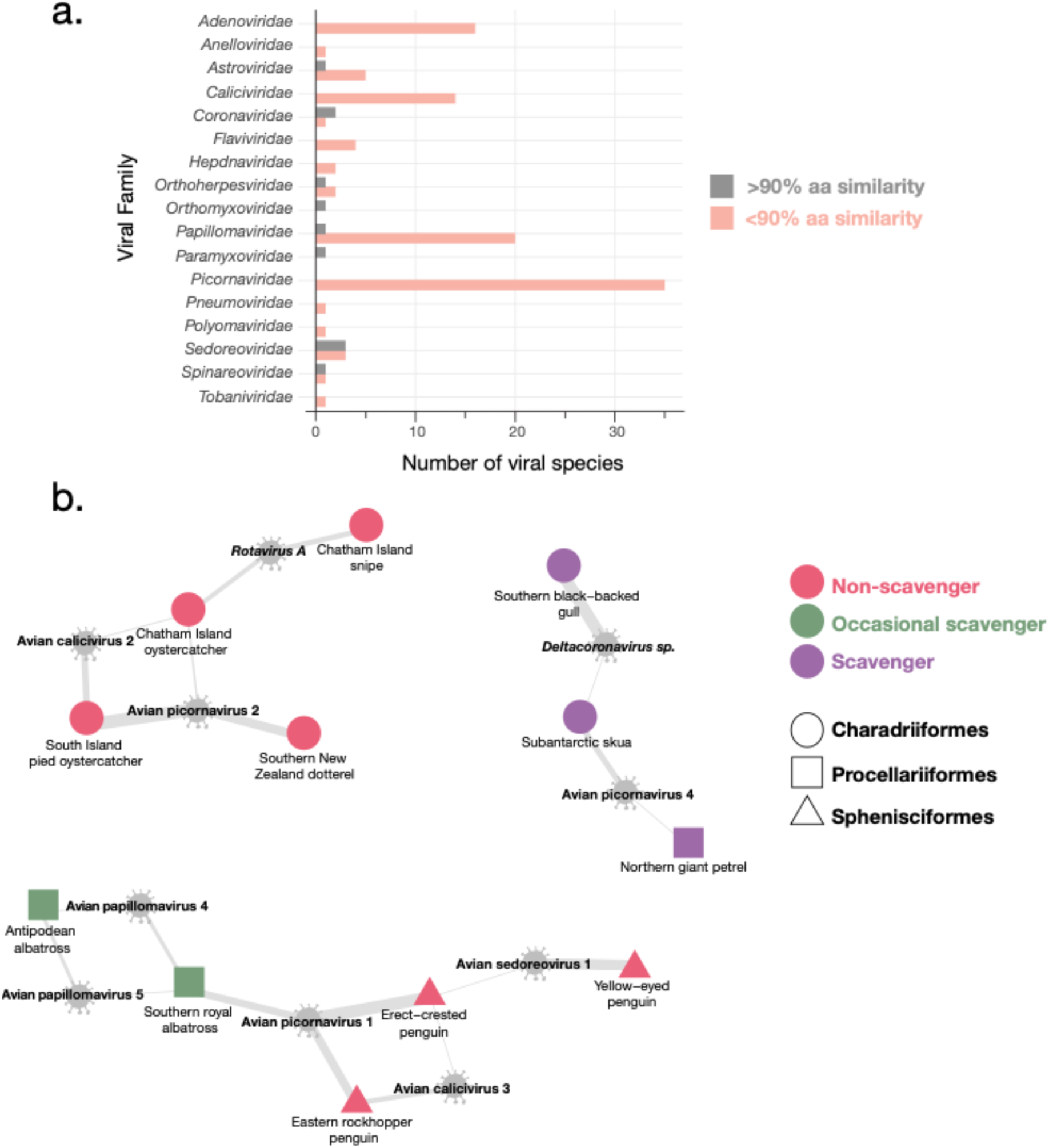
Viral species diversity and sharing among avian hosts. (a) The number of viral species identified within each viral family across all libraries. Viral species were separated into those sharing <90% amino acid similarity (pink) or ≥90% similarity (grey) with their closest BLASTp hit. (b) Bipartite networks illustrating viral species–host associations. Nodes represent viral species or avian hosts (node shape indicates avian order; node colour indicates scavenging status), and edges are weighted by relative abundance (RPM).

Although no highly pathogenic avian influenza viruses were detected, we identified the complete genome of a low-pathogenic H1N9 avian influenza virus, denoted A/Red knot/Firth of Thames/1/2024 (H1N9), in red knots sampled from the Firth of Thames on mainland New Zealand(16). Phylogenetic analysis of the hemagglutinin (HA) segment revealed close clustering (>90% nucleotide similarity) with H1 viruses previously sampled in Oceania and Asia(16). In addition, we discovered a highly divergent tobanivirus in an Auckland Island teal (*Anas aucklandica*) from Adams Island in the subantarctic region (Figure 3). To our knowledge, all previously described members of the *Tobaniviridae* have only been identified in mammals, reptiles, and fish. The divergent nature of this virus suggests it may represent the first avian tobanivirus reported to date.

To better resolve host associations and patterns of viral sharing we further examined the data at the viral species level. The viral families harbouring the greatest diversity of viral species were the *Picornaviridae*, *Caliciviridae*, *Papillomaviridae*, and *Adenoviridae* (Figure 4a). Most of the viruses identified (91%) exhibited <90% amino acid similarity to their closest BLASTp hits, indicating a high proportion of novel viral diversity. No novel viral species were detected within the *Paramyxoviridae* and *Orthomyxoviridae*, where only previously described avian viruses were identified.

We next explored host–virus associations to understand the prevalence and potential drivers of viral host-switching (Figure 4b). Most viral species were host-specific, with limited evidence of viral sharing across bird species. Where sharing did occur, it was largely among hosts with similar ecologies. For example, Avian picornavirus 2 was identified in Southern New Zealand dotterels (*Anarhynchus obscurus obscurus*), South Island pied oystercatchers (*Haematopus finschi*), and Chatham Island oystercatchers (*Haematopus chathamensis*), all Charadriiformes and non-scavenging birds. In addition, Avian papillomavirus 4 was detected in multiple albatross species classified as occasional scavengers. Viral sharing of Avian sedoreovirus 1 and Avian calicivirus 3 was also observed among penguin species (Sphenisciformes). In contrast, scavenging species, including subantarctic skuas, southern black-backed gulls, and northern giant petrels (*Macronectes halli*), harboured distinct viruses such as Deltacoronavirus sp. and Avian picornavirus 4 that were not shared with non-scavengers or occasional scavengers. Hence, viral sharing among bird species was limited, but when it did occur it tended to follow strong host-specific associations, often within closely related or ecologically connected avian species. Despite this, there was one notable exception, with Avian picornavirus 1 being identified in both species of Sphenisciformes (eastern rockhopper penguin (*Eudyptes filholi*) and erect-crested penguin (*Eudyptes sclateri*)) and Procellariiformes (southern royal albatross (*Diomedea epomophora*)) (Figure 4b).

To determine whether scavenging behaviour is phylogenetically structured, we tested the association between bird order and scavenging status. This is important because scavenging strongly influences virus exposure, particularly from other species or environmental sources such as carrion. A contingency table of bird order versus scavenging category (non-scavenger, occasional scavenger, scavenger) was analysed using Fisher’s exact test, which revealed a highly significant association (p <0.0001), such that scavenging is clustered within certain avian lineages. In our data set, scavenging was observed mainly in Charadriiformes (33% scavengers) and Procellariiformes (∼17% scavengers, ∼36% occasional scavengers), while Anseriformes and Sphenisciformes were entirely non-scavenging. As scavenging was non-randomly distributed across bird orders, differences in viral communities could otherwise reflect feeding behaviour rather than species identity or phylogeny. This justified including foraging category as an independent variable in downstream analyses of virome diversity and composition.

To assess the influence of ecological factors on viral community composition, we examined both alpha and beta diversity using Shannon indices and NMDS ordination (Figure 5). Alpha diversity differed significantly across foraging categories (Kruskal–Wallis χ² = 8.73, df = 2, p = 0.0127), with post hoc Dunn tests showing that scavengers had significantly higher Shannon diversity (mean = 0.25) than non-scavengers (mean = 0.13; p = 0.0095), while occasional scavengers (mean = 0.11) did not differ significantly from either group (Figure 5a).

**Figure 5.**
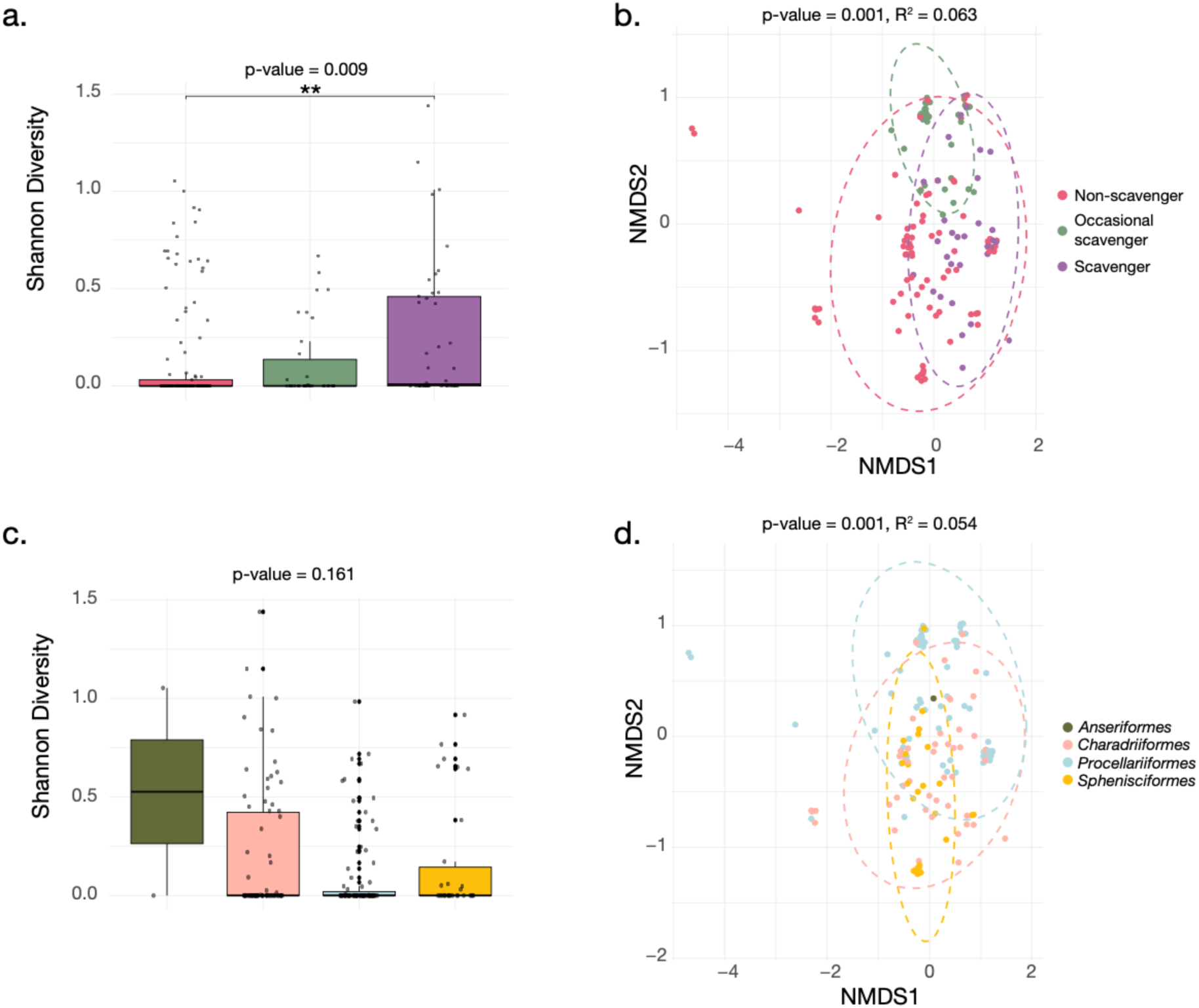
Alpha and beta diversity of avian viromes. (a) Boxplot of Shannon diversity (alpha diversity) across foraging status, with colours representing foraging behaviour. (b) NMDS plot of beta diversity coloured by foraging behaviour. (c) Boxplot of Shannon diversity across avian orders, with colours representing bird orders. (d) NMDS plot of beta diversity coloured by avian order. Significance values (p-values) and effect sizes (R²) are indicated above each panel where applicable.

**Figure 6.**
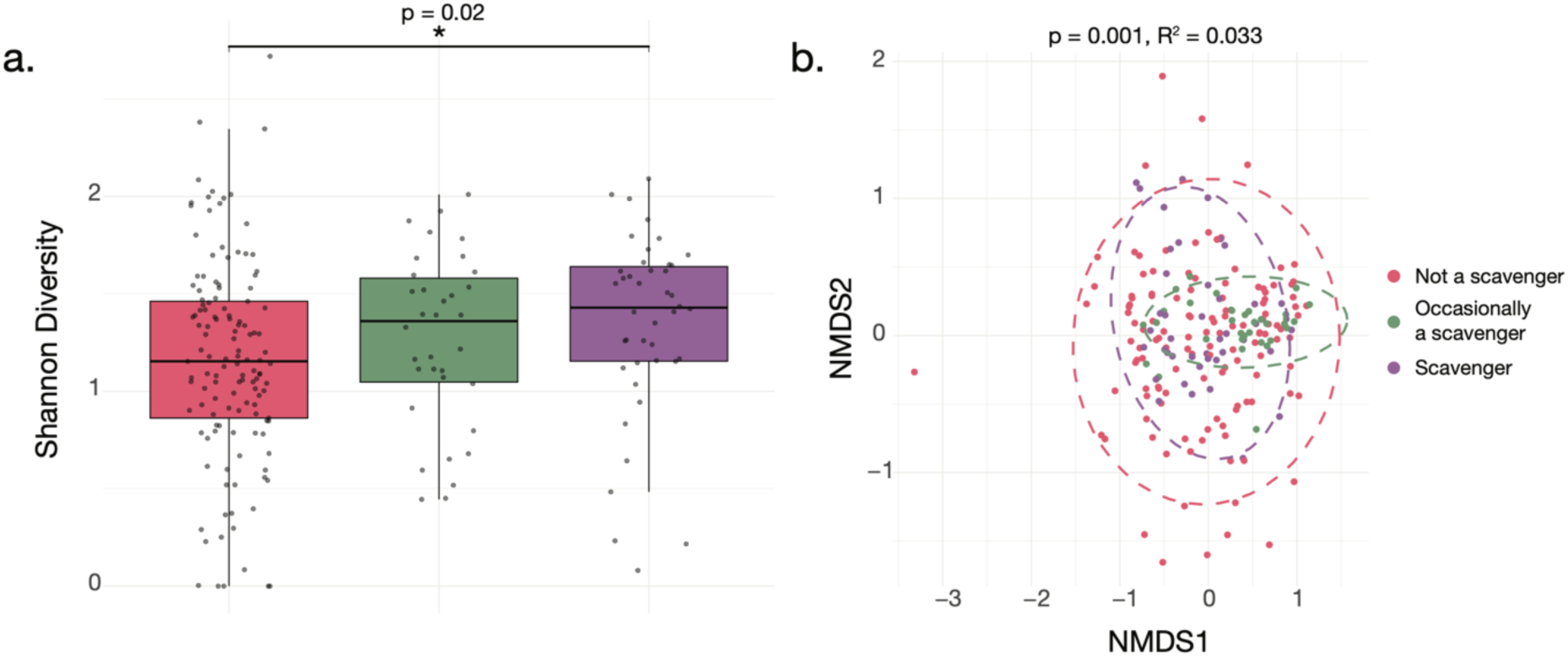
Alpha and beta diversity of viromes (non-avian viruses). (a) Boxplot of Shannon diversity (alpha diversity) across scavenging status, with colours representing scavenging categories. (b) NMDS plot of beta diversity coloured by scavenging status. Significance values (p-values) and effect sizes (R²) are indicated above each panel where applicable.

PERMANOVA on Bray–Curtis distances revealed a significant association between foraging behaviour and viral community composition (R² = 0.063, p = 0.001) (Figure 5b). Tests for homogeneity of dispersion (betadisper) indicated significant differences among foraging groups (p < 0.001), particularly between non-scavengers and scavengers, and between non-scavengers and occasional scavengers, suggesting that some of the observed differences in community composition may reflect variation in within-group dispersion as well as shifts in centroids. Nevertheless, even when accounting for the other factors (bird order (R² = 0.029, p = 0.007), geographic location (R² = 0.029, p = 0.006), sample type (R² = 0.017, p = 0.001), migration status (R² = 0.017, p = 0.065)) foraging status explained the largest proportion of variance (R² = 0.042, p = 0.001), highlighting its prominent role in structuring avian virome composition.

Indicator species analysis identified that some viral families were significantly associated with foraging behaviour. Specifically, the *Adenoviridae* (p = 0.005), *Astroviridae* (p = 0.014), and *Coronaviridae* (p = 0.019) were significantly associated with scavenging birds, suggesting these viruses may be more prevalent or abundant in species that feed on carrion. The *Papillomaviridae* (p = 0.004) were associated with both scavengers and occasional scavengers, indicating some viral overlap between these groups.

When considering bird taxonomy, no significant differences in Shannon diversity were detected, implying that overall levels of viral diversity did not vary substantially among different avian orders (Figure 5c). However, beta diversity analyses showed a statistically significant difference across bird orders (PERMANOVA, p = 0.001, R² = 0.054), indicating that while the overall diversity within each order was comparable, the specific composition of viral communities differed according to host phylogeny (Figure 5d).

To determine whether the influence of foraging behaviour on viral community composition extended beyond avian viruses, we repeated alpha and beta diversity analyses on non-avian viruses detected which spanned a total of 169 viral families (i.e. those associated with diet and the environment). Alpha diversity, measured by Shannon index, differed significantly among scavenging categories (Kruskal–Wallis χ² = 7.66, df = 2, p = 0.02), with scavengers showing the highest diversity (mean = 1.34 ± 0.07), non-scavengers the lowest (mean = 1.16 ± 0.05), and occasional scavengers intermediate (mean = 1.27 ± 0.08). Post hoc Dunn tests indicated that the significant difference was driven by scavengers versus non-scavengers (adjusted p = 0.02), while no significant differences were observed between occasional scavengers and either of the other groups. Beta diversity analyses using NMDS ordination also supported these patterns. PERMANOVA on Bray–Curtis distances confirmed that foraging behaviour significantly influenced viral community composition across non-avian viruses (R² = 0.033, p = 0.001). When accounting for additional ecological factors, bird order, sampling region, migration status and sample type, in a marginal-effects PERMANOVA, foraging behaviour remained a significant factor (R² = 0.020, p = 0.008), indicating a robust but small independent effect even in the context of multiple covariates. Other factors such as sample type (R² = 0.042, p = 0.001), bird order (R² = 0.028, p = 0.001), and sampling region (R² = 0.028, p = 0.002) were also significant, while migration status was not (R² = 0.012, p = 0.09).

Together, these results indicate that scavenging status influences virome diversity and composition not only among avian viruses but also when considering the total viral community, with scavengers consistently exhibiting higher alpha diversity and distinct viral community composition compared to non-scavengers.

Given the impact of scavenging on virus composition, we next examined the avian metatranscriptomes for the presence of mammalian-associated viruses (Supplementary Figure 1). Strikingly, across all libraries, we detected 12 mammalian viruses, including four that shared >90% amino acid similarity to previously classified viruses: *Hedgehog hepatovirus*, *Rabbit hemorrhagic disease virus 2*, *California sea lion astrovirus 5*, and *California sea lion astrovirus 8* which were identified in the metatranscriptomic libraries of southern black-backed gulls and subantarctic skuas. Ten of these 12 viruses were found in scavenging bird species, whereas only two were identified in non-scavenging birds, and the latter two viruses were sufficiently phylogenetically divergent that their true host origin remains uncertain. The predominance of mammalian viruses in scavengers is consistent with their feeding behaviour, which provides increased opportunities for acquiring viruses from carrion or mammalian by-products. Importantly, we also identified host-derived sequencing reads corresponding to hedgehogs, rabbits, and sea lions in all but one of the libraries in which their respective viruses were detected, with abundances ranging from 6-25 RPM, again consistent with scavenging behaviour (Supplementary Table 3). This co-detection strongly supports the detection of mammalian viruses as dietary or environmental acquisitions, rather than true infections of birds.

## Discussion

We performed the first large-scale metatranscriptomic survey of wild birds across New Zealand’s main migratory flyway sites and subantarctic islands. Our results reveal how viruses are transmitted among migratory and sedentary aquatic bird species across New Zealand, a key hub along major migratory flyways, at a time of heightened global concern over the spread of HPAI H5N1. In total we identified 118 avian viruses, of which 107 were putative novel species, demonstrating that New Zealand’s avian virosphere is vastly underexplored. In total 16 of the identified viruses shared close genetic relatives (>90% amino acid similarity) with avian viruses sampled from Australia, Antarctica, Japan, the Falkland Islands, Russia, China, and the United Arab Emirates, indicating that New Zealand’s avifauna is not completely isolated and shares viral lineages with multiple regions, although the mechanisms of introduction, whether via migratory species, vagrants, or other ecological processes, remain unclear.

To investigate the ecological and evolutionary drivers of avian virome composition, we tested the effects of host phylogeny, migratory behaviour, geographic location, and foraging behaviour on influencing virome diversity and composition. Surprisingly, migratory behaviour had a non-significant effect on beta diversity when accounting for other factors, indicating that migration alone did not significantly shape viral community composition in this dataset. Despite this non-significant result, there were examples within our data where migratory birds were likely bringing viruses in from overseas to New Zealand. For example, we identified that bar-tailed godwits (*Limosa lapponica*) harboured the same *Gammacoronavirus sp.* as previously detected in eastern spot-billed ducks (*Anas poecilorhyncha/zonorhyncha*) in Russia, and that subantarctic skuas harboured *Deltacoronavirus sp.* which was previously found in southern black-backed gulls in South Shetland Islands, Antarctica.

While viral diversity within each avian order was similar, community composition differed according to host phylogeny, indicating that likely phylogenetic barriers to transmission were shaping virome diversity and structure. Indeed, phylogenetic analysis revealed that viruses from hosts within the same avian order generally clustered together, reflecting host-specific associations, as did viruses sampled from the same geographic region. This trend was particularly evident within the *Papillomaviridae*, within which multiple viruses were detected across all four avian orders and all sampling regions but phylogenetically clustered according to both host order and location. Avian picornavirus 1 was the only notable exception, occurring in both Sphenisciformes and Procellariiformes sampled across different Subantarctic islands, indicating that viral sharing across host orders can occur when species occupy overlapping environments. Importantly, however, while avian order and geographic location significantly influenced beta diversity, the proportion of explained variation was lower than that associated with foraging behaviour, which is itself phylogenetically structured across bird lineages.

Foraging behaviour accounted for the largest proportion of variance in beta diversity (R² = 0.042), slightly higher than that explained by bird order (R² = 0.029), geographic location (R² = 0.029), and sample type (R² = 0.017). This suggests that foraging plays a relatively more prominent role in structuring avian viromes, but the differences among these factors are modest, highlighting that multiple ecological and phylogenetic factors jointly shape viral community composition. Scavengers, through their feeding habits, are more likely to encounter a diverse array of viruses from multiple species and environmental sources. This suggests that some of the viral diversity observed in scavenging birds may result from exposure to common scavenged sources rather than direct transmission among the birds themselves, although some bird-to-bird transmission cannot be ruled out. Similarly, snowy sheathbills (*Chionis albus*), opportunistic scavengers, were used as sentinel species for viral surveillance in Antarctica, where they were found to harbour a complex virome comprising avian, marine mammal, and human viruses(39). Foraging behaviour likely enhances exposure to a wide array of viruses, making scavengers important sentinels for emerging pathogens, and emphasising the need to integrate ecological and behavioural data when assessing virome dynamics and disease emergence risk.

In addition to avian viruses, we identified 12 mammalian-associated viruses. Notably, *Hedgehog hepatovirus*, identified in southern black-backed gulls, shared >99% amino acid sequence similarity and >87% nucleotide identity with *Hedgehog hepatovirus* previously reported in European hedgehogs (*Erinaceus europaeus*) sampled in Germany(40). That hedgehog reads were also present in the avian library (with an abundance of 6 reads per million) confirmed a dietary origin, indicating these detections likely reflected ingestion rather than active infection. This finding highlights the value of scavengers as sentinels of viral diversity across multiple species and ecosystems, providing a unique window into the movement of viruses through wildlife food webs. Given that southern black-backed gulls are largely resident within New Zealand and do not migrate internationally(41), this finding is unlikely to reflect recent viral importation. Instead, it suggests that the virus may have persisted within New Zealand’s introduced hedgehog populations since their introduction from Europe in the 19th century (42), maintaining high sequence conservation over time.

Broader temporal and geographic sampling, phylogenetic dating analyses, and targeted surveillance of migratory bird species and hedgehogs will be necessary to fully evaluate this hypothesis.

Other notable mammalian viruses include two sea lion astroviruses identified in subantarctic skuas. These astroviruses had >90% amino acid similarity with astroviruses previously reported in California sea lions (*Zalophus californianus*) sampled in California, USA(43). One showed close similarity (88% amino acid identity) to a California sea lion astrovirus that was also recently detected in New Zealand sea lions (*Phocarctos hookeri*), suggesting that their presence in skuas may reflect local acquisition rather than long-distance introduction(44). This finding remains important, as it highlights viral sharing among marine and avian hosts, and how migratory or scavenging behaviours, such as those of Subantarctic skuas which travel up to 6,000 km and frequently scavenge on marine mammal carcasses and intertidal resources, could facilitate ecological interfaces conducive to viral spillover (45). Indeed, repeated spillovers of HPAI H5N1 to marine mammals(46), and similar transmission pathways could theoretically facilitate the introduction of novel viruses to New Zealand. While the typically rapid onset of disease from viruses such as HPAI H5N1 may make long-distance transmission biologically unlikely in some cases(1), observations of avian migration demonstrate that birds can disseminate even highly pathogenic viruses across regions(1,19).

Importantly, HPAI H5N1 was not detected in any samples collected during the 2023–2024 sampling period, providing reassurance for biosecurity, conservation, and industry stakeholders. However, the threat of H5N1 introduction to New Zealand remains and is increasing, particularly since HPAI H5N1 has now spread to Indian Ocean archipelagos, demonstrating the continued potential for transboundary movement(19). Detection of a low-pathogenic avian influenza (LPAI) A/H1N9 virus in red knots demonstrates that migratory birds continue to act as reservoirs for diverse influenza viral strains(16). Notably, a recent outbreak of HPAI H7N6 in a poultry farm in New Zealand was most closely related to LPAI previously detected in mallard ducks (*Anas platyrhynchos*) in New Zealand in 2005, indicating that this virus likely emerged following local spillover of LPAI from wild birds to poultry(47). No closely related LPAI strains have been detected since 2015 in wild birds in New Zealand, suggesting that despite ongoing surveillance, gaps remain in our ability to monitor the diversity and dynamics of avian viruses in New Zealand’s wild bird populations and highlights the potential for undetected viruses to spill over into domestic poultry.

Together, these findings emphasise the importance of sustained surveillance and proactive biosecurity measures, and the need to integrate ecological, migratory, and virological data to inform risk assessments at regional, national, and global scales.

The identification of a divergent tobanivirus from the Auckland Island teal may represent the first putative avian member of the *Tobaniviridae*, broadening the known host range of this RNA virus family. This virus shares approximately 48% amino acid similarity with *Goat torovirus* yet forms a distinct lineage that does not cluster with established mammalian, reptilian, or piscine tobaniviruses. Until now, members of the *Tobaniviridae* have been found exclusively in mammals, reptiles, and fish. Auckland Island teal are flightless and endemic to the subantarctic Auckland Islands, having evolved in isolation from mainland New Zealand after diverging from their flighted relatives, the grey teal (*Anas gracilis*), chestnut teal (*Anas castanea*), and brown teal (*Anas chlorotis*), under relaxed selection pressures in a predator-free environment(48). Because Auckland Island teal primarily consume small marine invertebrates and terrestrial arthropods(49) and do not consume vertebrates, the identification of this novel tobanivirus is unlikely to reflect dietary contamination, suggesting a genuine avian host association. Under a model of virus-host co-divergence(50) it might have been expected that the novel avian tobanivirus would share common ancestry with reptilian tobaniviruses(50). Instead, the novel tobanivirus identified here shares common ancestry with mammalian toroviruses, indicating that tobaniviruses may have undergone complex host-switching events during their evolutionary history. Collectively, this finding expands the known evolutionary breadth of *Tobaniviridae* and highlights the potential of isolated island species like the Auckland Island teal to reveal deep and previously unrecognised viral lineages.

Overall, this study provides a baseline for the study of aquatic avian viromes in New Zealand, including sampling avian species from subantarctic territories and offshore islands. This work provides a reference for assessing future changes in avian virome composition, such as those resulting from incursions of HPAI H5N1. Importantly, our findings emphasise the role of foraging in shaping virome diversity, since scavenging birds are exposed to a wide range of viruses from multiple hosts and environments, making them key sentinels for the detection of emerging pathogens. These findings have direct relevance for wildlife conservation, particularly for New Zealand’s vulnerable endemic species that may be at risk from introduced viruses, and reinforces New Zealand’s value as a sentinel site for monitoring avian viral spread along global migratory flyways. Longitudinal monitoring will be critical for future investigations to capture seasonal and annual variation in viral diversity, while expanding sampling across additional species, geographic regions, and ecological niches will improve understanding of virus distribution. Integrating host ecology and movement data will further elucidate the drivers of viral transmission and evolution.

## Data availability

Viral sequences have been submitted to GenBank under the accession numbers [pending] while raw sequencing reads are available under BioProject [pending]. Original tree files for phylogenetic trees presented in Figure 3 can be found at https://github.com/stephwaller/NZ-Avian-Virome-2023-2024.

## Acknowledgements

We acknowledge Kaitiaki Rōpū ki Murihiku, Whenua Hou Komiti, and Kāi Tahu for allowing us to work on Southern Taonga. Successful sample collection in the subantarctic was possible thanks to Steve Kafka and the crew of the Evohe enabling safe passage to and from the islands. We also thank the Quarantine Store of the Murihiku Office of the Department of Conservation for ensuring the continued pest-free status of these unique islands. We would also like to acknowledge Environment Canterbury, Adrian Riegen, Johannes Chambon, Olivia Janes, Ela Hunt, Zoe Stone, Kaitlyn Hamilton, Jim Croawell, Bryan Annandale, Claudia Mischler, Rose Collen, Bruce Harrison, Harrison Talarico, Guy McDonald, Kevin Carter, Jim Fyfe, Hollie McGovern, Julia Reid and Brendon Clarke for contributing to bird sampling.

## Supplementary information

**Supplementary Table 1. Summary of sample pooling strategy and associated metadata for bird swab RNA samples.**

**Supplementary Table 2. Summary of avian and mammalian viruses identified from the metatranscriptomic libraries of avian samples.**

**Supplementary Table 3. Summary of mammalian host reads detected in the metatranscriptomic libraries of avian samples in which mammalian viruses were found.**

**Supplementary Figure 1. Phylogenetic trees of mammalian viruses.** Maximum likelihood phylogenetic trees were estimated from representative viral transcripts containing the RdRp gene from four viral families: *Astroviridae*, *Sedoreoviridae*, *Picornaviridae*, and *Caliciviridae*. Viruses identified in this study are shown in bold. Coloured dots beside viral names denote the scavenging status of the swabbed birds. Branch lengths represent the number of amino acid substitutions per site, and all trees are midpoint-rooted. Nodes with ultrafast bootstrap support values >70% are marked with an asterisk. Major viral genera within each family are highlighted.

