## Supplementary figures and images for "Foraging ecology drives viral community structure in New Zealand’s aquatic birds"

### Supplementary Figure 1

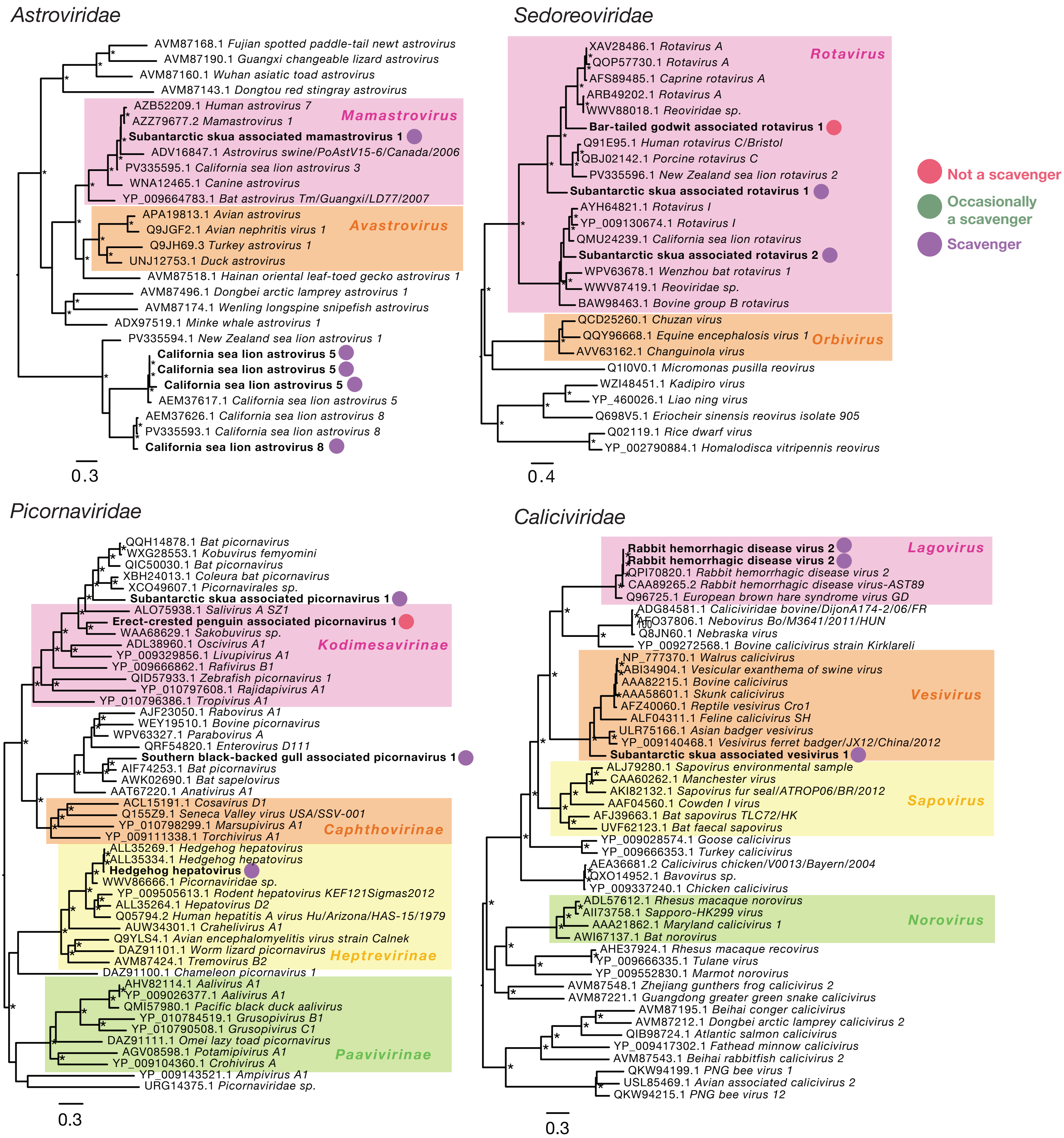
